# FastRecomb: Fast inference of genetic recombination rates in biobank scale data

**DOI:** 10.1101/2023.01.09.523304

**Authors:** Ardalan Naseri, William Yue, Shaojie Zhang, Degui Zhi

## Abstract

While rates of recombination events across the genome (genetic maps) are fundamental to genetic research, the majority of current studies only use one standard map. There is evidence suggesting population differences in genetic maps, and thus estimating population-specific maps are of interest. While the recent availability of biobank-scale data offers such opportunities, current methods are not efficient at leveraging very large sample sizes. The most accurate methods are still linkage-disequilibrium (LD)-based methods that are only tractable for a few hundred samples. In this work, we propose a fast and memory-efficient method for estimating genetic maps from population genotyping data. Our method, FastRecomb, leverages the efficient positional Burrows-Wheeler transform (PBWT) data structure for counting IBD segment boundaries as potential recombination events. We used PBWT blocks to avoid redundant counting of pairwise matches. Moreover, we used a panel smoothing technique to reduce the noise from errors and recent mutations. Using simulation, we found that FastRecomb achieves state-of-the-art performance at 10k resolution, in terms of correlation coefficients between the estimated map and the ground truth. This is mainly due to the fact that FastRecomb can effectively take advantage of large panels comprising more than hundreds of thousands of haplotypes. At the same time, other methods lack the efficiency to handle such data. We believe further refinement of FastRecomb would deliver more accurate genetic maps for the genetics community.

## 1 Introduction

A genetic map for a population or a species contains the locations of genetic markers or variant sites in relation to one another based on the probability of recombination, rather than a physical location along each chromosome. An accurate genetic map, which is an estimation of the recombination rates along a chromosome, serves as the foundation for genetic studies like gene mapping, population genetics, and genealogical studies. Given that recombination rates differ between populations, the estimation of population-specific genetic maps is crucial for advancing genetic research, particularly in diverse populations.

The genetic map is measured in centimorgans (cM), with each cM representing a 1% risk that two markers on the same chromosome would drift apart as a result of a recombination event during meiosis. In the human genome, one centimorgan roughly equates to 1 million base pairs (Mbps). The recombination rates may considerably vary within 1 Mbps, and the average of 1 cm equal to 1 Mbps may not hold at fine-scale (high) resolutions (see Figure 1). Some regions may also have significantly different recombination rates than the average.

**Fig. 1.**
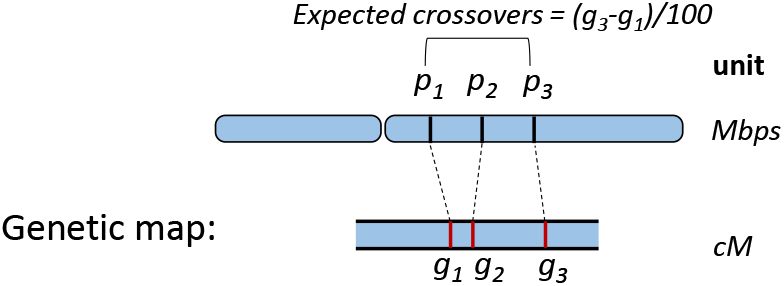
An example of a genetic map for a chromosome. The expected average number of intervening crossovers in a generation within (*p*_1_, *p*_2_) and (*p*_1_, *p*_3_) are (*g*2 – *g*1)% and (*g*3 – *g*1)%, respectively.

The traditional approach to infer the recombination rates is to use genotype data from a large number of parent-offspring pairs to capture an adequate number of meiotic crossover events [1, 2]. Fine-scale pedigree-based recombination rates from deCODE [2] are widely used. However, it is increasingly recognized that recombination rates vary from population to population [3]. Collecting a large number of parent-offspring pairs can be a practical bottleneck for most populations. An alternative approach is to use a single human sperm cell referred to as sperm-typing [4, 5]. A semen sample represents a significant portion of the meiotic crossover events because it contains hundreds of millions of sperm. Sperm-typing can predict a person’s unique recombination rate. However, at high resolution the rates may not always remain consistent with other individuals within the population. The sperm-typing is also a time-intensive and money-consuming process.

Another approach involves using population samples [6–11]. Recombination event signals in population samples are dispersed among the individuals. Not all recombination event signals, however, are correlated with the genetic map because they could have been brought by prehistoric crossovers. An early method based on population samples, LDhat, uses linkage-disequilibrium (LD) patterns for fitting a Bayesian model via MCMC [8]. While the results of LDhat are noteworthy, the limitations related to the computational tractability issues are significant as its capacity is restrained to a maximum of several hundred haplotypes. It is anticipated that the methods that can leverage large samples may achieve superior performance.

Recently, IBDrecomb [12] was developed to leverage the recent development of fast Identity-by-Descent (IBD) segment calling methods. IBD segments are identical DNA fragments that are inherited from a common ancestor. Under the assumption that the IBD segment boundaries were caused by recombination events, IBDrecomb counts the IBD segment boundaries and generates a map iteratively using the normalized counts of the IBD segments. However, IBDrecomb is not adequately efficient as it requires outputting all pairwise IBD segments, which is not conducive for biobank-scale cohorts. Here, we introduce a novel approach that efficiently identifies potential recombination breakpoints in very large cohorts using positional Burrows-Wheeler transform (PBWT). Our method is the first to directly use PBWT to estimate recombination rate. To enable accurate recombination rate estimation, we have the following methodological innovations and contributions: 1) Our method bypasses calling pairwise IBDs (e.g. IBDrecomb) to achieve the needed efficiency crucial for large cohorts. 2) The crossovers are counted by minor allele counts within individual PBWT blocks which is efficient while avoiding overcounting. 3) To avoid fragmenting PBWT blocks due to genotyping errors and recent mutations, we leveraged P-smoother. Our results confirmed the effectiveness of P-smoother.

## 2 Methods

### 2.1 Preliminaries

#### Positional Burrows-Wheeler Transform (PBWT)

The positional Burrows-Wheeler transform [13] facilitates an efficient approach for finding haplotype matches and also compression of haplotypes in large biobank-scale cohorts. The underlying idea of PBWT is to store the haplotype sequences based on their reversed prefix order. Following Durbins’s notation [13], we define a panel of haplotype sequences *X*, where *X* is a two-dimensional matrix. *X_k_* represents the values of haplotypes at the site *k* and *X* = [*X*_0_, *X*_1_,…*X*_*N*−1_], where *N* denotes the number of sites. *X_k_* is an array with *M* entries where *M* denotes the number of haplotype sequences. We also assume the entries of the array *X_k_* are binary.

#### Prefix array and PBWT matrix

The sequence indices sorted at each site *k* are referred to as the positional prefix array *a_k_*. PBWT matrix *y* stores the values of haplotype sequences in the reversed prefix order at each site. If divergence values and the haplotype sequences *X* are stored, there will be no need to store the PBWT matrix and the values at a site *k* for each haplotype *i* can be queries (*y_k_*[*i*] = *X_k_*[*a_k_*[*i*]]).

#### Divergence array

Divergence array *d_k_* at each variant site *k* for every haplotype stores the starting position of the match between the haplotype with its preceding haplotype sequence in the reversed sorted order up to the site *k* – 1. The divergence value keeps track of the starting site index of the longest match for each haplotype. We refer to the value of the divergence array for each haplotype as its divergence value. *d_k_* is utilized to both identify a long match with a length ≥ *L* and determine the starting position of the match.

#### Haplotype matching blocks for efficient identification of long matches

PBWT facilitates an efficient approach to enumerate all pairwise haplotype matches longer than a given length. While Durbin’s algorithm outputs all pairwise matches in *O*(*NM* +*#matches*), a block-based approach [14–16] can enumerate all matching blocks without explicitly outputting all pairs in *O*(*NM*). By sorting the haplotype sequences based on their reversed prefix order, the longest match for each haplotype sequence is placed in the adjacent position. Moreover, all pairwise haplotype sequences at the site *k* that are identical for at least *L* sites from *k* are separated by a haplotype sequence *j* with the condition *d_k_*[*j*] > *k* – *L* [13]. We define a *L*-block at a site *k* as a set of haplotype indices that share long matches with each other ending at site *k* with a minimum length of L. All L-blocks at any site *k* may be efficiently scanned by consensus PBWT algorithms [14–16].

#### PBWT-smoothing for reducing mismatches due to errors and mutations

The original PBWT scans the haplotype sequences starting from the first site. The divergence and positional prefix arrays are calculated at each site and the matches starting from the previous sites can be enumerated at each site. PBWT cannot tolerate mismatches in long matches. As a result, the long matches harboring mismatches will be discarded or reported partially depending on the minimum cutoff length. The bi-directional PBWT data structures [17], on the other hand, provide an efficient approach to tolerate possible mismatches in the middle of long matching blocks.

In order to tolerate genotyping errors, the haplotype panel is smoothed using bi-directional PBWT. The pre-processing step alternates the alleles that are different in the middle of matching blocks of haplotypes. The smoothing procedure (P-smoother) only alternates the minor alleles if the minor allele frequency is below a certain threshold (with the minor allele frequency threshold equal to 5% by default) in the matching blocks. This allows our method to be highly error-tolerant and maintain accuracy even when subjected to genotyping errors.

### 2.2 Inferring recombination rates

Similar to pedigree-based [2] and population-based inference methods [12], we use an iterative approach to count crossover events of a certain type at each bin (or window) across the genome. To estimate the recombination rates efficiently, the type of crossover events should be chosen that can be efficiently counted and unbiased across the genome. Instead of counting all IBD segment boundaries, we count the boundaries of diverging haplotypes across all matching blocks in each bin. Haplotypes *diverging* at site *k* are defined as the haplotypes that are matching with at least one other haplotype until the site *k* – 1, and the match between the haplotype and other haplotypes in the block terminates at the site *k*. As we consider haplotypes in a block are split into two clusters, one having the major allele and the other the minor allele at site *k*. We can call the haplotypes carrying the minor allele *diverges* from the block with haplotypes carrying the major alleles.

Given a haplotype panel comprising *N* sites, the recombination rates are calculated for each window in terms of physical distances (default *w* = 5000). We assume that the total genetic length in cM is known. We iterate over the sites and the number of minor alleles for haplotypes within any *L*-block are counted. The number of total recombination events in each window is simply the sum of all minor allele counts from the sites in the window *i*. Our approach avoids enumerating all pairwise haplotype matches at each site. Please note that the time complexity of enumeration of all pairwise matches, especially for very short segments could be theoretically quadratic. Theoretically, the *#matches* could be *O*(*M*^2^) for very short matches. Our method only iterates over the haplotypes at each site and computes the number of minor alleles in each block. In addition, the divergence values for the haplotypes with the minor alleles are then considered to update the value for the preceding windows containing their divergence values. The overall time complexity of our method is *O*(*NM*), where N denotes the number of sites and M denotes the number of haplotypes.

Figure 2 shows a simple example of a haplotype panel sorted by their reversed prefix order at sites *k*. The haplotypes marked with a red cross at the site *k* are being considered in the calculation of recombination events. For each *L*-block, the haplotypes with minor alleles are considered and the allele count for the window overlapping with their divergence value is updated. Each block is likely derived from a different genealogical branch. The ancestral accumulation of recombination events is therefore controlled by considering *L*-haplotype blocks. Algorithm 1 describes the procedure of counting possible recombination events at each window using PBWT arrays. The array *g*[*i*] contains the genetic location of the site *i*. *L* denotes the minimum genetic length for a match. As we iterate over the haplotypes in the reversed prefix order, the condition *g*[*d_k_*[*j*]] > *g*[*k*] – *L* triggers the enumeration of possible recombination events. The array *count* stores the total count of recombination events (minor allele counts) for each window. The array *pos* contains the genomic (physical) position of the sites.

**Fig. 2.**
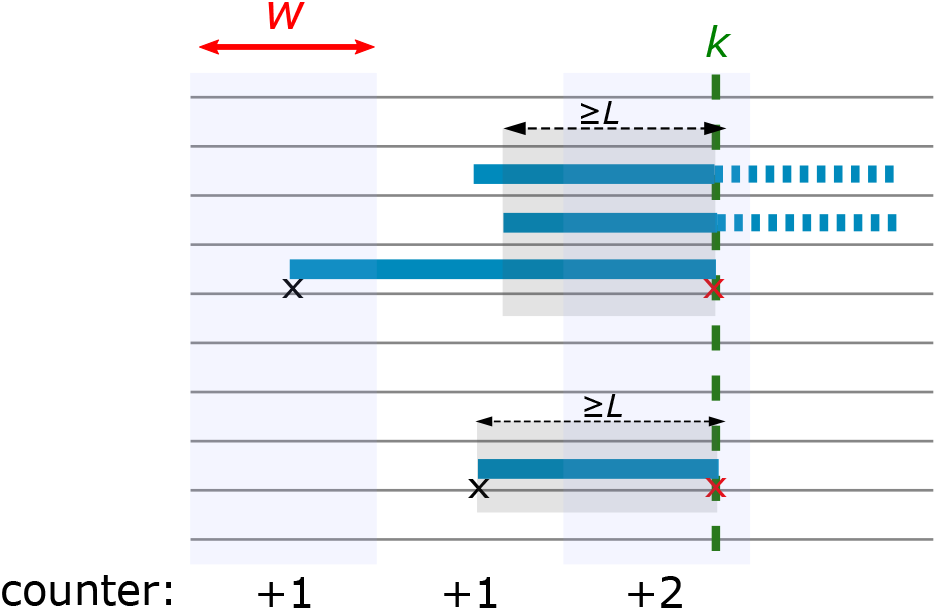
A sketch of the FastRecomb algorithm for counting recombination events at the variant site *k*. The array counter stores the sum of events for all sites within each window with the length *w*. Gray horizontal lines represent haplotypes in the panel. Haplotypes are sorted based on their reversed prefix order at site *k*. Two L-blocks are highlighted in light gray. The diverging haplotypes (red ‘x’s) in each L-block are counted and the windows containing the starting of their longest match are updated.

#### Algorithm 1 countAltAlleles

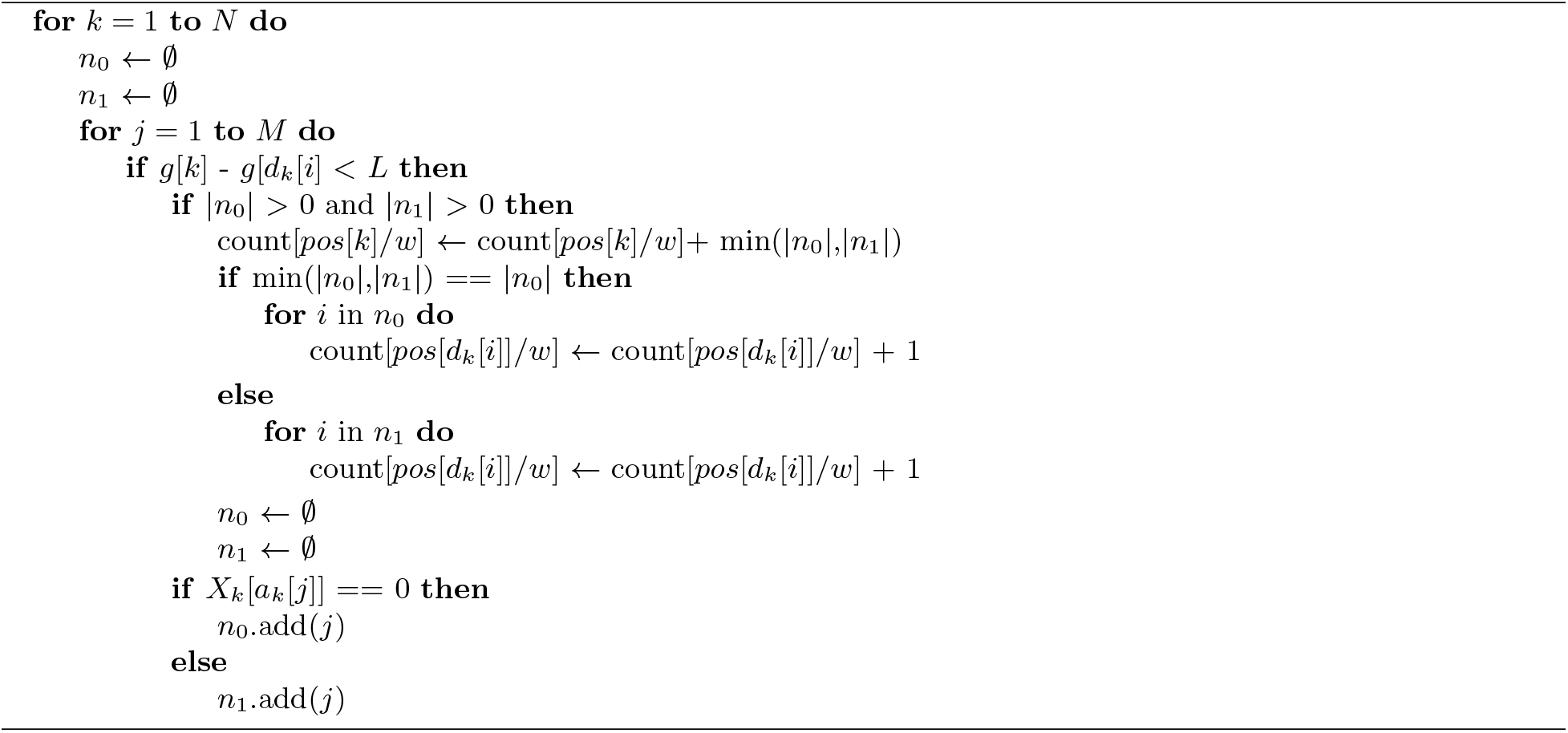

The recombination rate for each window *i* is calculated by the formula:

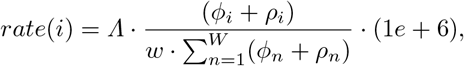

where *Λ* denotes the total chromosome length in cM, and *w* is the window size in terms of physical distance. *ϕ_i_* and *ρ_i_* represent the number of minor allele counts among the haplotype blocks that end in the *i*-window and start, respectively. We start with a simple assumption of a constant recombination rate across the chromosome (1 cM ≈ 1 Mbps). In the first iteration, the matches are considered by assuming the minimum length is in Mbps. In each iteration, for each window, the average between the current calculated rate from the previous iteration is considered.

## 3 Results

### 3.1 Simulation

We simulated 1 million haplotypes of European and African ancestry using deCODE genetic map [1]. OutO-fAfrica_2T12 model in *stdpopsim* [18] was used to simulate haplotypes of chromosome 20 with the command line: `stdpopsim HomSap -c chr20 -d OutOfAfrica_2T12 -g DeCodeSexAveraged_GRCh36 500000 500000` *stdpopsim* uses the coalescent simulator *msprime* [19] as its simulator engine. The variant sites with an allele frequency of less than 0.05 were filtered out. The total number of variant sites after the MAF filter was 66,546. We also inserted different genotyping errors into the simulated panel.

### 3.2 Evaluation of estimated rates

To evaluate the performance of our method, FastRecomb ^3^, the Pearson correlation coefficients for different resolutions were calculated. The highest resolution was set to 10,000 which was the resolution of the genetic map in deCODE36. We compared the performance of FastRecomb with IBDrecomb and LDHat. The current implementation of FastRecomb does not treat the end regions differently as was done in IBDrecomb. As a result, the values for the end regions may not be optimal. Here, we focus on the inferred recombination rates in the mid-region (excluding 5 Mbps from both sides of the chromosome). For FastRecomb, we first ran `P-smoother` [20] with parameters *L′* = 20, *W′* = 20, *g* = 1, *MAF* = 5% and then ran `FastRecomb` with parameters *L* = 66.3, *d* = 0.5, *w* = 5000, *r* = 5. *r* denotes the number of iterations and *d* denotes the minimum target length in cM. For LDhat, we ran the `interval` method with a block penalty of 5 and for 22.5 million iterations with a sample being taken every 15k iterations. For IBDrecomb, we ran `refined-ibd` [21] with a minimum LOD score of 1 and a minimum IBD segment length of 0.3 cM. We then ran `merge-ibd-segments` with a gap of 0.6 cM and discord of 1. Due to the resource-intensive and run time requirements of LDhat and IBDrecomb, we used only 192 and 5k haplotypes, respectively. For LDhat, 192 haplotypes were the largest number of haplotypes for which a pre-computed likelihood lookup table was available. For IBDrecomb, 5k haplotypes were the size of the simulated data in their study. We also tried running `refined-ibd` on 100k haplotypes and the program had not terminated after a month of running.

Figure 3 shows the correlation coefficients of the three methods for the mid-region (excluding 5 Mbps from both ends of the chromosome). No genotyping errors were added to the haplotype panel. FastRecomb performs better than other methods for the mid-region in different resolutions. For 500k resolution, all the methods achieve a high correlation coefficient close to 1. We also analyzed the five highest recombination rate locations in deCODE (Chr20) to examine how the hotspots are replicated. We compared the top five recombination rates inferred by different tools to deCODE using different distance cut-offs. Two from FastRecomb and IBDrecomb, and one from LDhat regions were within 10k of the five deCODE hotspots. FastRecomb, IBDrecomb, and LDhat scored 3, 2, and 2 for a distance cut-off of 400k, respectively. Running on an 8-core 2.10 GHz Intel Xeon E5-2620 v4, LDhat (`interval` + `stat`) took 2.39 CPU hours, IBDrecomb (`refined-ibd` + `IBDrecomb`) took 83.6 CPU hours (approximately 3.25 wall-clock hours), and FastRecomb (`P-smoother` + `FastRecomb`) took 13.2 CPU hours. Please note that only 192 haplotypes were used for LDhat and 5k for IBDrecomb. FastRecomb’s time complexity is linear to the sample size, hence it is possible to run the program with one million haplotypes without using extensive resources. The maximum resident set size (peak memory) for FastRecomb was only ~ 77 MB. The peak memory values for LDhat and IBDrecomb were ~ 781 MB and ~ 810 MB, respectively.

**Fig. 3.**
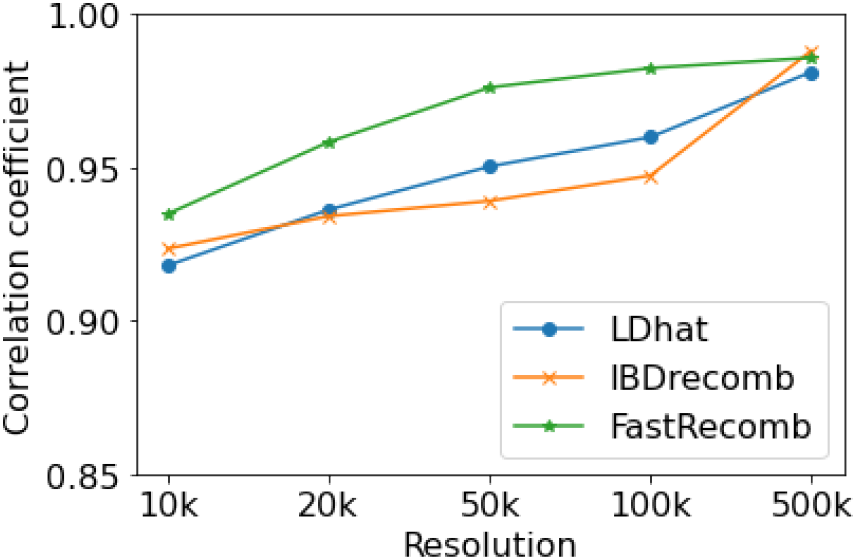
Pearson correlation coefficients between the inferred recombination rates and the ground truth.

### 3.3 Robustness against genotyping errors

Genotyping errors in real datasets are more dominant than mutations. It is almost impossible to ignore the genotyping error rates in practice. We evaluated the performance of FastRecomb, IBDrecomb, and LDhat using different genotyping error rates. We implanted error rates of 0.05, 0.1, and 0.2%. The errors were randomly inserted for each haplotype. For example, to simulate an error rate of 0.1%, we randomly selected 0.1% of the variant sites for each haplotype and altered the alleles. Figure 4 shows the correlation coefficients for the three methods in mid-region using 0.05% (a), 0.1% (b) and 0.2% (c) error rates. The correlation coefficients of FastRecomb are not affected by increasing the error rates (see Fig 4). For IBDrecomb, however, additional genotyping errors decrease the correlation coefficient values. LDhat, similar to FastRecomb, appears to be robust against error rates up to 0.2% for each haplotype. We repeated the experiment with the genotpying error of 0.1% for 10 different panels generated using varying seed values *s* = {1,…10} (see Figure 5).

**Fig. 4.**
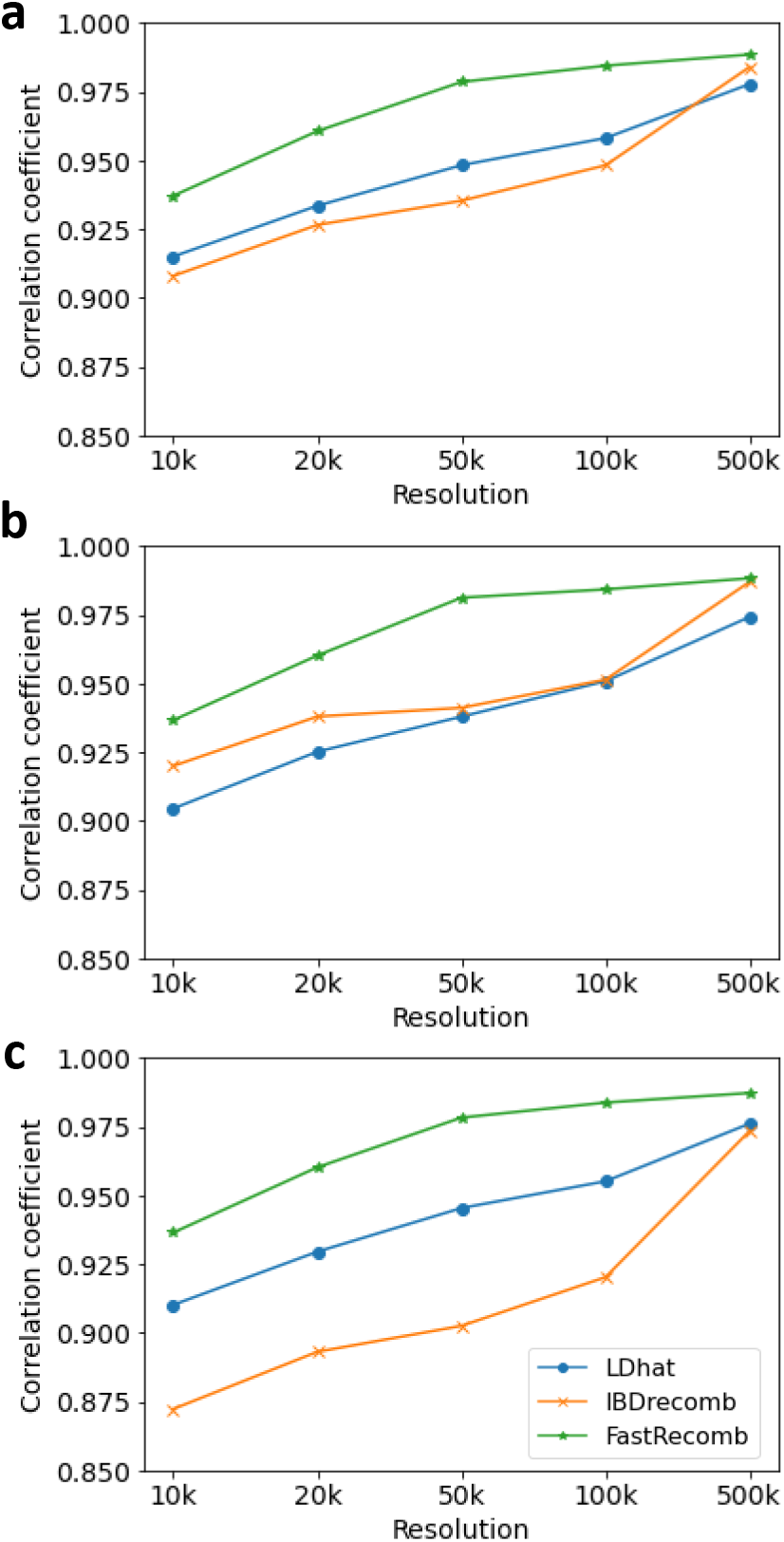
Pearson correlation coefficients in panels containing different error rates; 0.05% (a), 0.1% (b) and 0.2%(c).

**Fig. 5.**
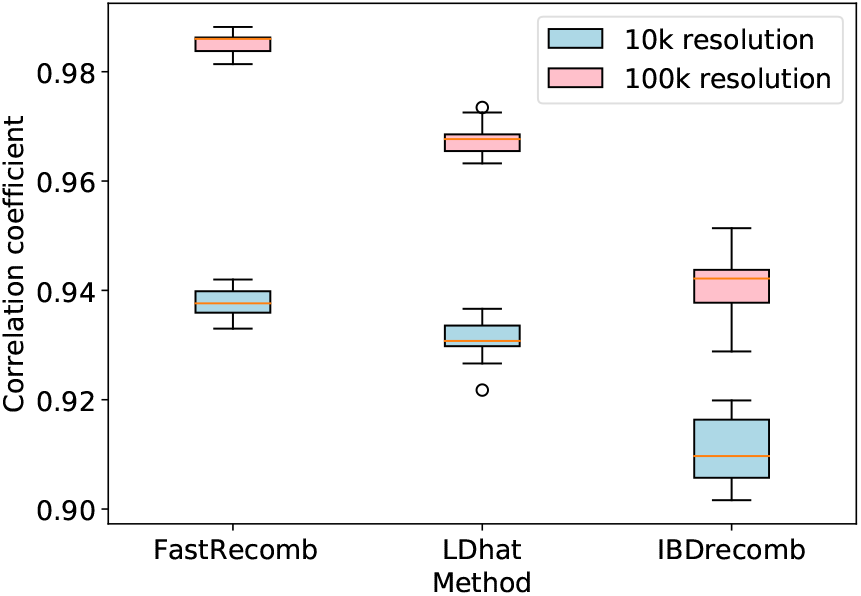
Performance of recombination rate estimation tools at 10k and 100k using 10 different panels with a genotyping error rate of 0.1%.

### 3.4 Performance growth with increasing sample size

FastRecomb performance depends on the number of samples from a population. Principally, FastRecomb facilitates the use of information that large-scale genetic data entail. To evaluate the performance of FastRecomb with increasing sample size, we extracted subpanels with 20k, 50k, 100k, 200k from the simulated 1 million haplotypes. Figure 6 illustrates the correlation coefficients for the mid-region using different sample sizes (from 20k to 1M haplotypes). The error rate per haplotype was set to 0.1%. The results of LDhat and IBDrecomb have been included as dotted and dashed lines, respectively. The sample sizes for LDhat and IBDrecomb were 192 and 5k. As shown in Figure 6, the correlation coefficients for FastRecomb increase with the increasing number of haplotypes. For smaller sample sizes (e.g. 20k) the coefficients from rates calculated by FastRecomb are lower than IBDrecomb and LDhat. With 100k haplotypes, the correlation coefficients in 50k resolution approach the values of IBDrecomb and LDhat. With 200k, the coefficients from FastRecomb are slightly higher than other methods in 50k resolution. With 1 million haplotypes, the correlation coefficients from rates calculated by FastRecomb exceed the values from LDhat and IBDrecomb in both 10k and 50k resolutions.

**Fig. 6.**
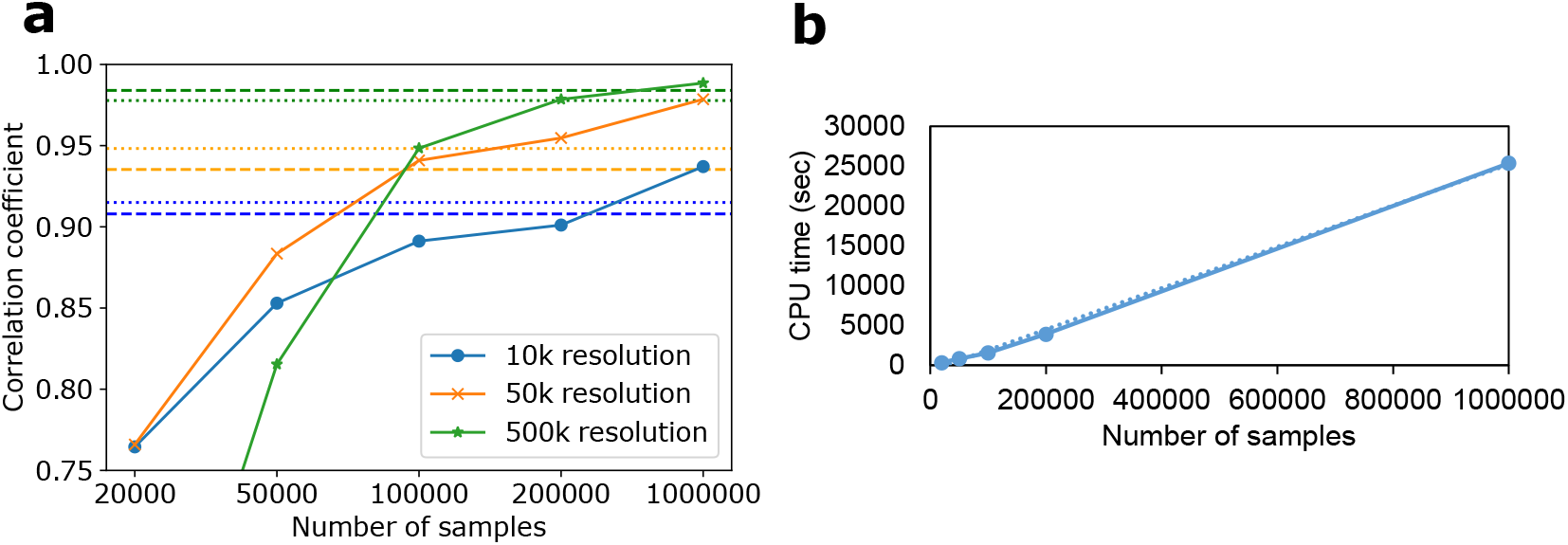
Effect of sample size on the performance of FastRecomb. Pearson correlation coefficient of FastRecomb improves with the increasing number of haplotypes (a). The running time increases linearly with the sample size (b). The error rate was set to 0.1%. Dashed and dotted lines represent the IBDrecomb and LDhat results, respectively.

### 3.5 Performance of FastRecomb without smoothing

The smoothing pre-processing step is crucial for panels with a high number of genotyping errors. We calculated the correlation coefficients of FastRecomb without the smoothing steps for different error rates (see Fig. 7). As shown in Figure 7, a low error rate (e.g. 0.05%) may not affect the results significantly. Higher error rates, however, could lead to a significant performance reduction.

**Fig. 7.**
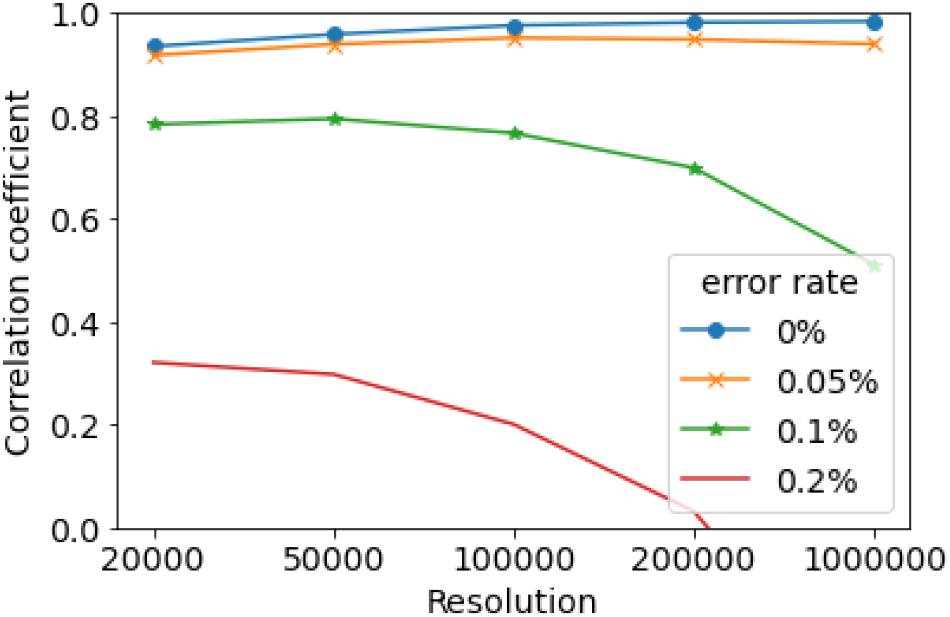
Pearson correlation coefficients of FastRecomb without smoothing the haplotype panel using different error rates.

Table 1 contains the correlation coefficients for the mid-region using smoothing and without smoothing. The genotyping error rate was set to 0%. The results show that the smoothing would not lower the correlation coefficients even if no genotyping error was expected.

**Table 1.** Correlation coefficients for mid-regions in a panel without genotyping errors.

|  | 10k res. | 20k res. | 50k res. | 100k res. | 500k res. |
| --- | --- | --- | --- | --- | --- |
| with smoothing | 0.9352 | 0.9588 | 0.9761 | 0.9818 | 0.9839 |
| w/o smoothing | 0.9348 | 0.9582 | 0.976 | 0.9824 | 0.9858 |

### 3.6 Performance of FastRecomb in end-regions

The coefficient values for end-regions in 10k and 500k resolutions from the panel with 0.1% error rates have been included in Table 2. LDhat shows a better performance in end-regions. For the end region 2-5 Mbps, the difference between LDhat and FastRecomb is less noticeable in 10k resolution. The current implementation of FastRecomb does not treat the end region differently. Hence, the correlation coefficients for the end-regions are not as high as those of the mid-region. IBDrecomb treats end-regions differently from the mid-region by a special procedure to compensate for the lower IBD boundary counts due to chromosomal ends. The special treatment by IBDrecomb improves the accuracy of the rates in the end-region, but, for LDhat, the correlation coefficients in the end-region are still higher.

**Table 2.** Pearson correlation coefficients for end-regions and mid-region for different recombination rate inference methods. The error rate was set to 0.1%. The end-regions contain 10 Mbps and the mid-region contains 47 Mbps of the chromosome.

|  | 10k resolution |  |  | 500k resolution |  |  |
| --- | --- | --- | --- | --- | --- | --- |
|  | <2 Mbps | 2-5 Mbps | mid-region | <2 Mbps | 2-5 Mbps | mid-region |
| LDhat | 0.9281 | 0.9520 | 0.9149 | 0.9962 | 0.9907 | 0.9777 |
| IBDrecomb | 0.8262 | 0.8799 | 0.9079 | 0.9960 | 0.9908 | 0.9840 |
| FastRecomb | 0.8321 | 0.9179 | 0.9370 | 0.8852 | 0.8855 | 0.9885 |

### 3.7 Recombination rate estimation in real data

We applied FastRecomb on four different subsets of UK Biobank data: 1) Asian or Asian British individuals, 2) Black or Black British individuals, 3) White individuals and 4) White (subset) individuals containing 7816 randomly selected samples. The ethnic background (Data-Field 21000) of only 8034 individuals was Black or Black British individuals. Therefore, we selected a subset of White individuals similar to that of Black or Black British individuals. The number of White (subset) individuals with available genotype data was slightly less than 8034. Approximately 1 CPU hour and 55 MB memory were used to estimate the recombination rates for the largest panel containing all White individuals within the UK Biobank data. Figure 8 illustrates the estimated rates for each subsets in chromosome 22. Table 3 contains the correlation coefficients and the number of individuals for different subsets of UK Biobank samples.

**Fig. 8.**
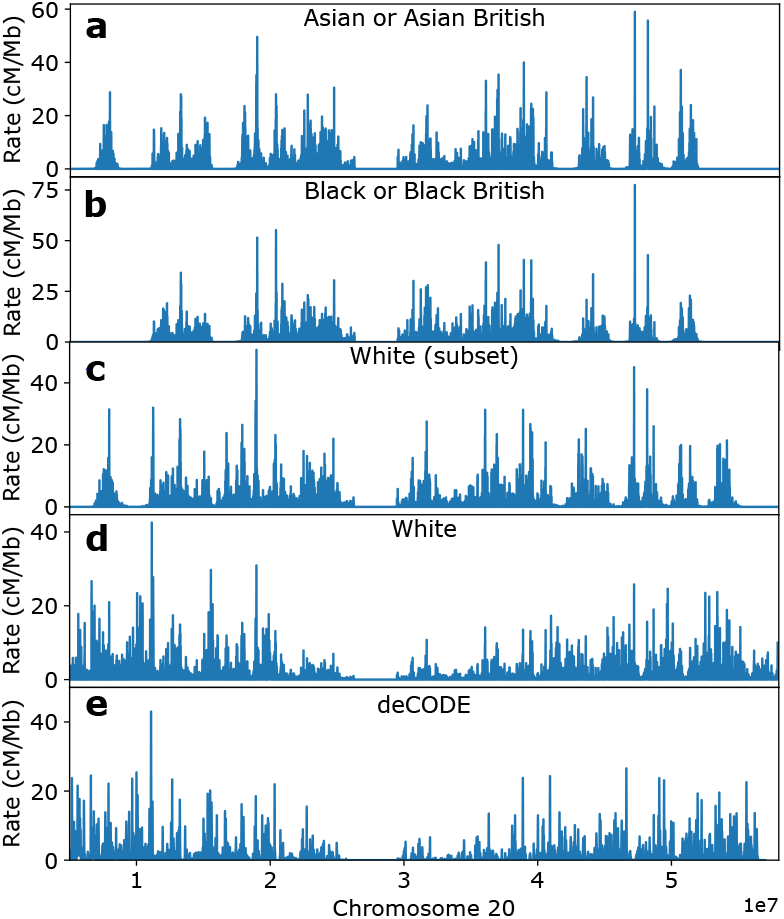
Estimated rates using different subsets of UK Biobank across the chromosome 20. (a) Asian or Asian British, (b) Black or Black British, (c) Subset of White individuals, (d) All White individuals. The rates from deCODE (sex averaged) are presented for comparison (e).

**Table 3.** Correlations between inferred rates in UK Biobank and deCODE map at 100k resolution.

| Ethnic background | Number of individuals | Correlation coefficient |
| --- | --- | --- |
| Black or Black British | 7618 | 0.26 |
| Asian or Asian British | 9375 | 0.33 |
| White (subset) | 7816 | 0.42 |
| White | 458677 | 0.79 |

Correlation with White (subset) was lower than White (all), confirming the higher power of larger sample size. Also, among populations with comparable sample sizes, the similar population has higher correlation, suggesting FastRecomb captured population-specific recombination maps. Note that even the highest correlation (0.79) between the estimated map from 458,677 (British) White individuals and the deCODE map are not as high as that from simulation (e.g., Table 1) This may be due to the fact that the genetic map between the Icelandic population (deCODE) may be different from the UK population, among other factors.

## 4 Discussion

The recombination rates inferred by FastRecomb can achieve the highest correlation coefficients given a large number of samples. The scalability of FastRecomb which enables the usage of biobank-scale data is perhaps the most significant reason for increased performance outcome. Additionally, the pre-processing step and the use of a shorter IBD length cut-off ensure robustness against genotyping errors, which would otherwise reduce the accuracy of IBD-based approaches like IBDrecomb. In our simulated data, panels with 100k individuals achieve comparable or slightly better performance compared to LDhat and IBDrecomb in the mid-region of the chromosome. With 500k individuals, the recombination rates inferred by FastRecomb are more accurate in the mid-region. In our experiments, we set the minimum length *L* for counting minor alleles in blocks of matching haplotypes to 0.5 cM. For smaller panels, the IBD coverage for 0.5 cM may not be sufficient for certain regions. As a result, the inferred recombination rates may not be representative of the underlying population. Smaller cut-off lengths (e.g. 0.1 or 0.2 cM) could result in more accurate correlation coefficients for small panels comprising only a few thousand haplotypes, but the correlation coefficients may not be necessarily better than the longer cut-off in large panels. This is due to the high probability of a match between two haplotypes for small cut-offs (e.g. 1 cM), especially if the marker density is not very high.

FastRecomb assumes that mismatches after a long match are due to recombination events where some mismatches could be due to recent mutations or genotyping errors. This problem can be addressed by smoothing the panel. However, we did not assess the biases by counting minor alleles, which could be further investigated in future work. In our experiments, we focused on the mid-region of the chromosome which entailed approximately 82% of the entire chromosome. In future work, we will experiment with special treatment for the end-regions similar to IBDrecomb while counting the IBD boundaries. Moreover, we limited evaluation of our method to array data. Error rates in sequencing data might be higher, especially for rare variants. As a result, the accuracy of FastRecomb will rely on the performance of the smoothing step. Thinning the data using minor allele frequencies is also a possible solution for panels with high error rates in rare variants.

## 5 Conclusions

In this work, we presented a new method to estimate the recombination rates in biobank-scale cohorts. A unique hallmark of the proposed method, FastRecomb, is its scalability. FastRecomb implicitly considers all matches between pairs of haplotypes while avoiding enumerating all possible pairs using PBWT blocks. As a result, the run time of FastRecomb grows linearly with the number of variant sites and the number of individuals. Also, FastRecomb avoids explicit outputting of IBD segments, a potential I/O bottleneck. These innovations enable FastRecomb to be easily applicable to panels with hundreds of thousands or even millions of haplotypes without requiring extensive resources.

Based on our experiments, the performance of FastRecomb can be higher than the state-of-the-art methods when the haplotype panel is large enough. Moreover, our method is robust against genotyping errors as the performance of FastRecomb was not affected by increasing the error rates from 0 to 0.2%.

In summary, FastRecomb unleashes the power of biobank-scale haplotype panels for estimating population-specific genetic maps. Given the fact that some human populations may have unique recombination hot-spots and their genetic map may differ, it is essential to estimate their unique genetic map for downstream analysis. As access to biobanks housing hundreds of thousands to millions of individuals increases, efficient methods such as FastRecomb may become critical for population-specific genetic map estimation.

## Acknowledgements

This work was supported by the National Institutes of Health grants R01HG010086, R56HG011509, and OT2OD002751.

## Footnotes

3 Source code is available at https://github.com/ZhiGroup/FastRecomb

